# Benchmarking Spectral Library Prediction Platforms for Neuropeptidomics Applications

**DOI:** 10.64898/2026.08.07.743122

**Authors:** Lauren Fields, Emily M. Hubecky, Kendra G. Selby, Lingjun Li

## Abstract

Data-independent acquisition (DIA) mass spectrometry has emerged as a powerful tool for neuropeptidomics, but its success relies heavily on the quality of spectral libraries used for peptide identification. There are inherent challenges to mass spectrometry analysis of crustacean neuropeptides, including the endogenous nature in which they are analyzed, extensive post-translational modification (PTM), and atypical fragmentation patterns. Thus, general-purpose proteomic spectral prediction tools may not perform optimally in the endogenous peptide domain. In this study, we benchmark four widely used spectral prediction platforms, Prosit, MS2PIP, AlphaPeptDeep, and UniSpec, to evaluate their performance in predicting the fragmentation of neuropeptides. Using an empirically derived spectral library from crustacean tissues as reference, we assess model compatibility, dot-product similarity, Pearson correlation, and DIA-based identifications across brain, sinus gland, and pericardial organ samples. Our results reveal that no single model comprehensively captures neuropeptide fragmentation characteristics. While UniSpec showed unexpected strengths due to its inclusion of neutral loss ions, AlphaPeptDeep demonstrated the highest spectral similarity, and MS2PIP and Prosit outperformed in DIA-NN identifications. We further highlight the critical impact of neutral loss fragments, present in over 50% of empirical spectra, and emphasize the need for hybrid spectral libraries that integrate complementary strengths across models. This work provides a foundational framework for optimizing spectral library selection in neuropeptidomics and underscores the importance of model-specific biases when analyzing structurally diverse endogenous peptides.

## Introduction

Spectral libraries serve as the foundational interpretive framework for peptide identification in data-independent acquisition (DIA) workflows, where fragment ions from multiple co-isolated precursors are acquired simultaneously and must be accurately matched to known fragmentation patterns. Unlike traditional sequence-database searching, which relies on theoretical fragment generation, spectral libraries encode the empirical or predicted fragmentation behavior of peptides, capturing sequence-specific intensities, PTM-driven fragmentation pathways, and characteristic neutral-loss patterns that cannot be inferred from sequence alone.[1] In neuropeptidomic analyses, where peptides are non-tryptic, structurally heterogeneous, and prone to producing b-ion–dominant and neutral-loss–rich spectra, spectral libraries are not merely supportive, they are indispensable.[2, 3] They provide the ground truth that DIA algorithms use to disentangle multiplexed spectra, reduce false assignments, and recover peptides that would be missed by purely theoretical models.[4] Consequently, accuracy and completeness of a spectral library directly determine the depth, precision, and reproducibility of DIA-based neuropeptide identification.

We have previously applied spectral libraries, complied from data-dependent acquisition (DDA) experiments, to facilitate neuropeptide identification in DIA workflows.[4, 5] While these empirical libraries have enabled the identification of numerous endogenous peptides, they are not exempt from the limitations inherent to experimentally acquired resources including sparse sequence coverage, incomplete representation of low-abundance species such as neuropeptides, and the absence of novel or unexpected peptide products.[6] As predicted spectral libraries have become increasingly robust and are rapidly emerging as the dominant strategy in modern DIA analyses, it is now essential to critically evaluate how well current prediction tools perform for analytes whose fragmentation behaviors differ substantially from those of classical bottom-up proteomics.

This need is particularly pronounced in the context of crustacean neuropeptidomics, where our knowledge remains comparatively limited. Approximately 1,000 crustacean neuropeptides have been identified to date,[4, 5] but many exist only as transcriptomic predictions without empirical MS/MS validation and others appear as partial degradation products of larger prohormones. These truncated peptides cannot be dismissed, as short neuropeptide fragments often retain biological activity.[7] Yet, this small and heterogeneous peptide pool is insufficient to train or fine-tune deep learning based spectral predictors, which typically require hundreds of thousands to millions of annotated spectra for stable optimization.[8] As a result, independent model development or domain-specific retraining remains infeasible for neuropeptides, forcing reliance on existing prediction tools originally developed for well-behaved tryptic peptides.

Neuropeptides, particularly crustacean neuropeptides, present distinct analytical challenges that diverge sharply from the assumptions encoded in most prediction frameworks. Unlike proteolytic peptides, endogenous neuropeptides are non-tryptic, non-digested, and often feature irregular termini and extensive post-translational modifications.[4] These features produce MS/MS fragmentation patterns dominated not by y-ions, a canonical feature of tryptic peptide spectra,[9] but frequently by b-ions, which are far more prominent in many neuropeptide families.[10] Additionally, neuropeptide spectra often rely heavily on neutral loss ions, including water, ammonia, or modification-specific moieties, which frequently exceed the intensity of backbone fragments and serve as key diagnostic indicators.[11] Because spectral prediction models are largely trained on tryptic peptides with minimal neutral-loss contribution, they struggle to recapitulate the true fragmentation behavior of endogenous neuropeptides.

While many modern prediction frameworks allow for retraining or fine-tuning with specialized datasets, such options are inaccessible for analytes that lack sufficient empirical depth or for researchers without advanced computational expertise. With only ∼1,000 known crustacean neuropeptides, such retraining would be statistically irresponsible and technically unstable. Thus, it becomes necessary to evaluate existing prediction models directly, leveraging their inherent strengths while recognizing the biases introduced by their training domains.

Among the available prediction models, Prosit, one of the earliest deep-learning frameworks, includes features designed to broaden its applicability.[12] We were especially interested in the Prosit 2025 model, which incorporates additional fragment types that more closely align with the complex fragmentation patterns characteristic of neuropeptides,[13] as well as the Prosit 2024 model, which expanded PTM support to better accommodate the modification-rich nature of endogenous peptides. MS2PIP, a long-standing machine-learning model widely used in proteomics, provides consistent intensity predictions and integrates well with DIA tools such as DIA-NN.[14] Its training on primarily tryptic peptide datasets, however, shapes the types of fragmentation patterns it most accurately predicts.

Other models have adopted different strategies. UniSpec incorporates a wider array of fragment ions, including several neutral-loss species, providing an alternative representation of peptide fragmentation that may be informative for certain endogenous peptide classes.[15] Its overall coverage and prediction characteristics vary across peptides with different structural features. AlphaPeptDeep, a transformer-based deep-learning framework, offers considerable flexibility in modeling sequences and PTMs and is designed to generalize beyond traditional proteomic peptides.[16] As with all models, its predictions reflect the characteristics of its training data, which may influence how well it captures neutral-loss-rich or non-tryptic fragmentation events. Collectively, these tools represent a diverse set of prediction strategies, each bringing different modeling assumptions and fragment representations that may impact their performance on neuropeptides.

## Methods

### Dataset description

The spectral library used for initial benchmarking (*i.e.,* empirical library) was described elsewhere and acquired on a Thermo Exploris 480 mass spectrometer equipped with high-field asymmetric-waveform ion mobility spectrometry (FAIMS).[5] In brief, 12 male Jonah crabs, *Cancer borealis*, were anesthetized on ice and dissected, obtaining brain, sinus gland (SG), and pericardial organ (PO) tissues. Tissues were extracted in acidified methanol and fractionated via offline high-pressure liquid chromatography (HPLC) to improve sample resolution using a Kinetex® Core-Shell EVO 5 μm C18 column on a Waters Alliance 2695 Separations Module operating at 0.2 mL/min. Fractions were analyzed on an Orbitrap Exploris 480 Mass Spectrometer (Thermo Fisher Scientific, CA) coupled to a Vanquish Neo UPLC system (Thermo Fisher Scientific, CA) in DDA mode. To build the spectral library, raw files were converted to .mzML and .MS2 file formats via MSConvert[17] and RawConverter,[18] respectively. Database searching was conducted in EndoGenius with the following parameters: parent mass error tolerance, 20 ppm; fragment mass error tolerance, 0.02 Da; m/z range, 50-3000 m/z; minimum precursor intensity, 1000; maximum precursor charge, 8; maximum fragment charge, 4; maximum PTMs per peptide, 3; confident coverage threshold percent, 70%; EndoGenius Score threshold, 1000. PTMs included C-terminal amidation, oxidation of methionine, N-termini cyclization of glutamic acid and glutamine, and sulfation of tyrosine.

The DIA evaluation data were also previously published, and acquired with the same instrumental setup as the described DDA experiment.[5] Tissues from 5 Jonah crabs were prepared as before, and data was acquired in DIA mode, with a 3 s cycle time and 10 *m/z* isolation window and 1 *m/z* window overlap.

### General prediction parameters

All libraries were predicted with the same parameter components, where possible. For AlphaPeptDeep, specification of instrument is required; Q Exactive and Lumos models were independently surveyed, as these were the instruments available.[16] Instrument selection was also required in UniSpec; we selected Q Exactive and Q Exactive HF-X,[15] which most closely resemble the Orbitrap instrument utilized in the empirical library. A collision energy of 30 eV, and a fragmentation selection of HCD was applied, where required. Predictions were obtained via Koina API (Accessed April 26, 2025). All neuropeptides within an in-house database were queried at precursor charges 2 through 7 (Supplementary File 1). All PTMs represented in the empirical spectral library were queried wherever possible, though some were incompatible with particular models. In the same way, retention times were predicted via the Koina API.

### Library evaluation

Libraries were evaluated in DIA-NN (v. 1.8.1). Libraries were converted to .TSV for compatibility with DIA-NN and analyzed with the aforementioned modifications (*e.g*., C-terminal amidation, Met oxidation, pyro-Glu from E, pyro-Glu from Q, sulfation of Tyr) at a Q-value of 0.05. For endogenous peptide evaluation, *in silico* cuts at any residue were prohibited. All Python scripts used for prediction and evaluation are freely available at https://github.com/lingjunli-research/NeuropeptideSpectralLibraryPrediction.

## Results and Discussion

### Overview of models

The recent release of several platforms that integrate multiple spectral prediction models, such as Koina[13] and Oktoberfest[19], enable the simultaneous prediction of spectra without imparting the time and sample burdens that are imposed with more traditional, experimental-based spectral library curation. With these advances, it has become clear that systematic evaluation of model performance is necessary to determine which tools are appropriate for specific analytes. We therefore began by assessing our neuropeptide spectral library and its agreeability with several widely used models.

Neuropeptides display distinctive characteristics in mass spectrometry compared to tryptic proteomics workflows. Neuropeptides are routinely identified with precursor charges ranging from +2 to +8,[4] extending beyond the expected maximum tryptic peptide precursor charge (**Fig. S1A**). This is in part driven by their wide mass range (**Fig. S1B**); neuropeptides can be extremely short, (*e.g.,* biologically active tripeptides[20]) or extremely long (*e.g.*, 70+ residue crustacean hyperglycemic hormones[21]). These deviations from traditional proteomics workflows are further reinforced by the recent observation that endogenous peptides frequently display high b-ion abundance, a clear departure from the typical y-ions dominance of tryptic peptides.[10] Furthermore, mature neuropeptides commonly possess N-terminal cyclization of glutamate and glutamic acid (pyro-Glu) and C-terminal amidation, [22, 23] modifications that are relatively rare in proteomics and not routinely incorporated into prediction model training. These features underscore the need for preliminary evaluation before adopting general-purpose spectral prediction models for neuropeptidomics.

Compounding this challenge, neuropeptidomics researchers have limited ability to retrain prediction models due to the small size of available datasets. For example, a recent transfer-learning study successfully adapted a tryptic model for HLA peptides using ∼100,000 peptides for both training and testing.[16] While this demonstrates the power of model customization, the crustacean neuropeptidome contains only ∼1,000 annotated peptides,[24] two orders of magnitude fewer, restricting similar approaches. Moreover, despite speculated similarity between HLA peptides and neuropeptides, this relationship has not been rigorously assessed, and crustaceans lack an adaptive immune system,[25] making HLA-based data unlikely to represent crustacean endogenous peptides.

### Model comparison

We compared four intensity-prediction model families: Prosit, AlphaPeptDeep, MS2PIP, and UniSpec (**Table S1**). Prosit, one of the most influential models in proteomics, has undergone substantial evolution since its inception. The 2019 model was the least compatible with neuropeptides, supporting only tryptic peptides with fewer than 30 amino acids with no PTMs.[12] The 2020 model expanded training to non-tryptic peptides,[26] and the 2024 model added support for 21 PTMs; however peptides remain limited to 30 residues. Koina also offers an unpublished 2025 Prosit model with improved handling of multiply charged fragments.[13]

MS2PIP followed a similar trajectory, originally restricted to tryptic peptides containing less than 30 residues,[27, 28] but has since expanded to include models covering diverse instruments, fragmentation methods, and quantification schemes. Most relevant here, the Immuno-HCD model incorporates non-tryptic immunopeptides,[29] another class of endogenous peptides which are likewise not subjected to digestion prior to analysis. AlphaPeptDeep, comparatively, is trained on both tryptic and non-tryptic peptides with no strict length limitations and a limited PTM set.[16] UniSpec was the only model without non-tryptic training; however, it supports peptides up to 40 residues, precursor charges up to +8, and a wide variety of PTMs.[15]

We first assessed which models could successfully generate predictions for neuropeptides in our empirical spectral library (**Fig. 1A**). Across prediction models, UniSpec yielded the greatest number of predictions, followed by both the Immuno and HCD models of MS2PIP, and lastly AlphaPeptDeep and both the 2019 and 2020 models of Prosit each (**Fig. 1A**). Despite this presumed initial success, UniSpec, AlphaPeptDeep, and Prosit all showed poor compatibility with neuropeptides, with MS2PIP showing markedly better compatibility (**Fig 1A**). Though this may initially suggest greater success of MS2PIP to predict neuropeptide spectra, many of the compatible entries produced zero dot-product similarity to empirical spectra (**Fig. 1A**). These “N/A” outputs were most common in MS2PIP, with observations across all models.

**Figure 1:**
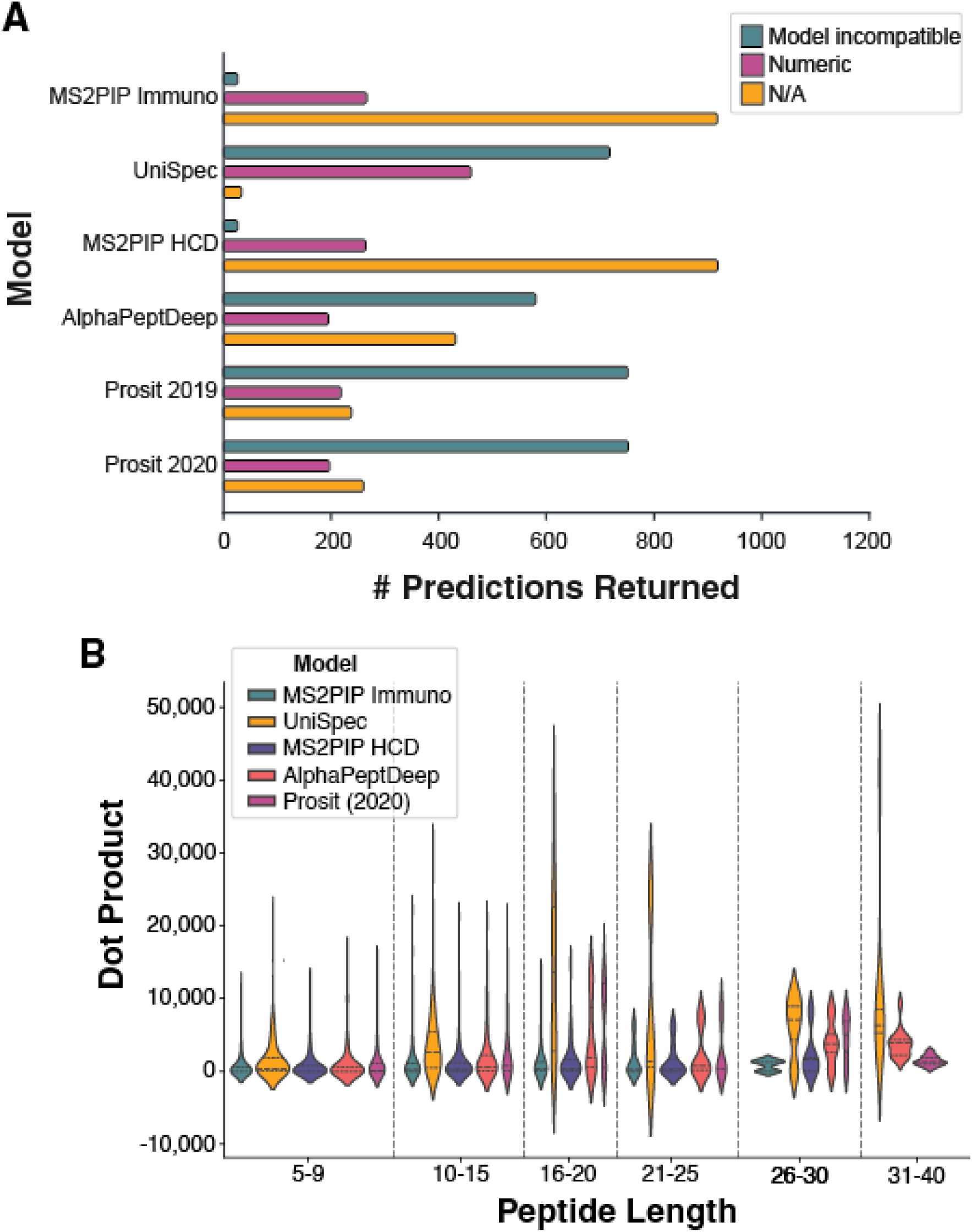
Results of each model in library prediction. **A)** Each model returned predictions, summarized by a numeric dot product (pink). Additionally, some returned zero peaks, denoted in yellow. Some models also were completely incompatible with the requested peptide predictions (blue bars) **B)** The length of the peptides predicted and their corresponding dot product distribution across each intensity model.

Unexpectedly, UniSpec produced the largest number of compatible predictions (**Fig. 1A**) and showed higher dot-product values across peptide-length (**Fig. 1B**), precursor-charge (**Fig. S2**), and PTM categories (**Fig. S3**). This was surprising given that UniSpec lacks non-tryptic training. These findings were of particular interest as they were a stark departure from the anticipation that an immunopeptide model, or a model trained on non-tryptic peptide, would be superior to this work. This presented the notion that each of the libraries are incredibly sensitive to their training datasets, and comprehensive evaluation is needed to determine the context in which a library best performs.

Given the variety of model versions, we next evaluated the sensitivity of prediction settings to dot-product similarity. Prosit 2019, 2020, and 2024 showed similar behavior while the 2025 model was substantially different and far less consistent (**Fig. S4A**). UniSpec models were highly consistent across all available instruments (*i.e.*, Thermo Fusion Lumos, Q Exactive, Q Exactive-HFX), while the generic model diverged more noticeably (**Fig. S4B**). In particular, agreement between Q Exactive and Q Exactive-HFX models was more pronounced than with the Fusion Lumos, reflecting instrumental similarities.[30–32] We also identified a slight difference between the MS2PIP HCD and Immuno HCD models (**Fig. S4C**), as well as from the AlphaPeptDeep Lumos and QE models (**Fig. S4D**), likely reflecting model improvements and instrumental differences.

We next evaluated the relationship between *m/z* tolerance to Pearson correlation coefficient (PCC), to determine the appropriate error margin for utilization of predicted libraries. Moreover, this evaluation offered the ability to profile the sensitivity of the models, wherein a lower *m/z* error tolerance should reveal a higher PCC. It would be expected that the “optimal” error margin would be between 0.01 and 0.02 *m/z*, consistent with high-resolution orbitrap instrumentation. In this evaluation, we realized that the MS2PIP HCD model, when at this optimal range, produces a PCC approximately equal to 0.75 (**Fig. S5A**), while the Immuno HCD model reveals a PCC closer to 1.0 (**Fig. S5B**). This emphasizes the sensitivity that is improved by the model tailored toward endogenous peptides. As anticipated, almost identical results can be observed for the AlphaPeptDeep Lumos (**Fig. S5C**) and Q Exactive (**Fig. S5D**) models; however, the PCC interestingly is consistent with values of 1.0 up to an error tolerance, or bin size, of 0.1 *m/z*. In a similar way, the UniSpec models for Lumos, QE, QE-HFX, and generic instrumentation (**Fig. S5E-H**, respectively) reveal nearly identical PCC values, though even at the lowest error margin considered, 0.01 *m/z*, a number of entries presented a PCC of −1, revealing negative correlation, and several entries congregating near 0, revealing no relationship. This highlights that, while the UniSpec model has advantages of producing a theoretical spectrum with potentially important neutral loss fragments, the model is not translational to neuropeptides at a global level. With Prosit, we see that the 2019 model (**Fig. S5I**) has a reasonable PCC cutoff at 0.1-0.02 *m/z*, while the 2020 model is more sensitive, with correlation controlled up to a threshold of 0.05 *m/z* (**Fig. S5J**), and the 2024 model yet even more sensitive to a threshold of 0.1 *m/z* (**Fig. S5K**). Deviating from its earlier counterparts, the Prosit 2025 model overall lacks correlation and proved to be insensitive at any threshold, with nearly even distributions of positive and negative correlation values even at the most restrictive error threshold (**Fig. S5L**). These findings validate the standard 0.02 *m/z* threshold, while determining that for the UniSpec and Prosit 2025 models, even restrictive tolerances do not represent correlation to endogenous neuropeptides.

We then compared the spectral libraries predicted by each model to each other and to empirical data. When initially surveying peptides with no modifications, a strong agreement between models is revealed (**Fig. S6**). This trend is disrupted upon introduction of PTMs, where performance of the models is more profound (**Fig. 2**). Of note, there is clear correlation of models generated by the same research groups, for example the UniSpec QE and QE-HFX model have an average PCC of 0.97;however, similarity across models was largely quite low. Despite this, we noted that the AlphaPeptDeep Lumos and QE peptides have a PCC of 1.00 and 0.99, respectively, with the empirical data. The Prosit 2019 and 2020 models likewise demonstrated high correlation with empirical data, with an average dot product of 0.93 and 0.99, respectively. When considering that several models had a large fraction of peptides incompatible with their model, this average was weighted against the known neuropeptidome, with each neuropeptide considered at several charge states. It is here that we see that the AlphaPeptDeep models appear to still be in strong agreement with the experimental spectra (**Fig. S7A**), and also the highest number of compatible peptide queries (**Fig. S7B**),hinting at the effectiveness of this model. Additionally, it is worth recognizing that the MS2PIP models evaluated, HCD 2021 and Immuno HCD had mean PCC values of 0.77 and 0.95 (**Fig. S7A**), respectively, despite having the same number of compatible peptides (**Fig. S7B**). This highlights that the model tuned for immunopeptides is substantially more effective at resembling expected spectra compared to the tryptic-only model.

**Figure 2:**
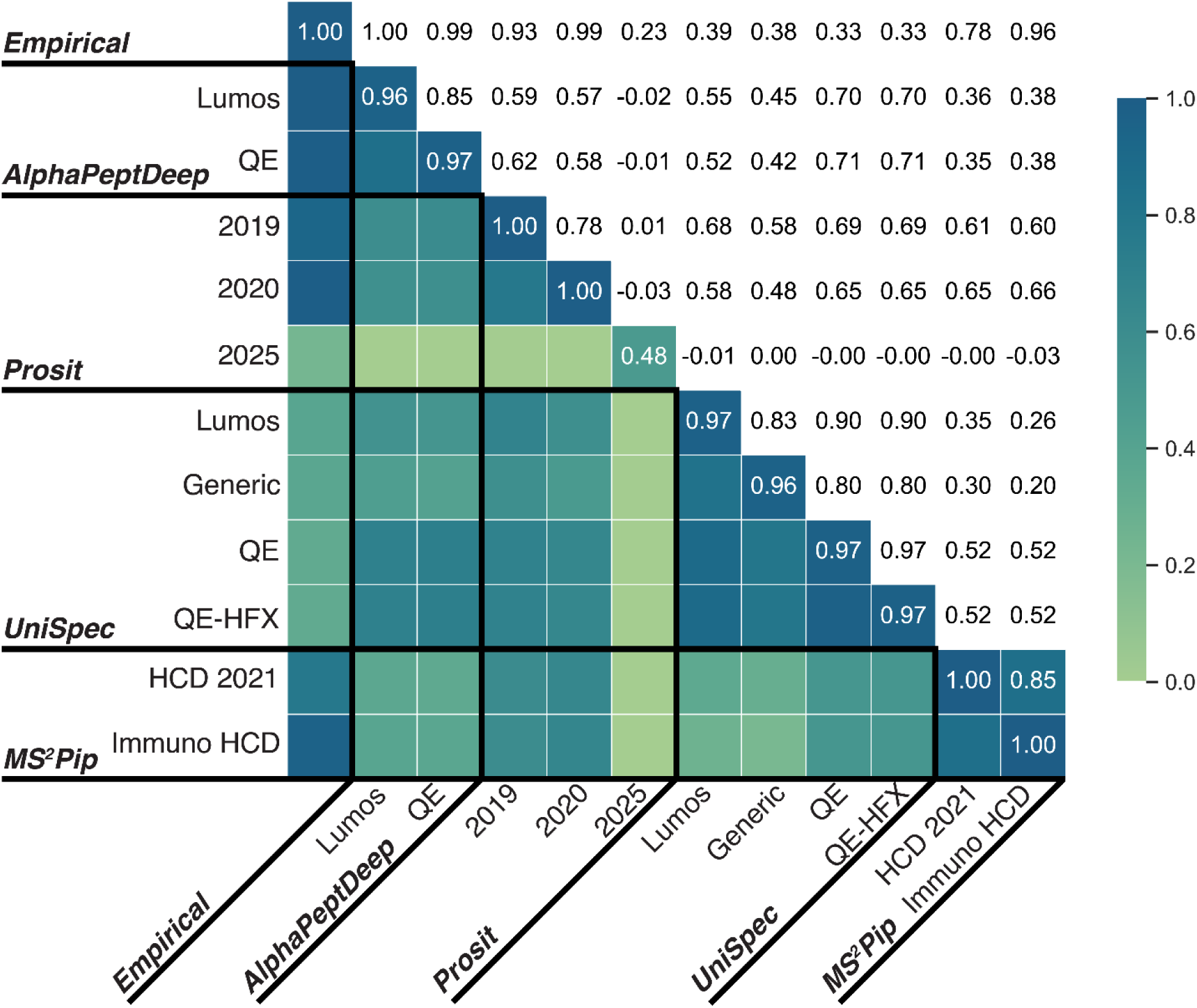
Average Pearson correlation coefficient (PCC) of each intensity model when compared to all other models.

### Retention Time Model Evaluation

For retention time (RT) prediction, Koina provides several models. In this work, we used AlphaPeptDeep’s Generic model, Chronologer, DeepLC, and the Prosit models from 2019 and 2024. For AlphaPeptDeep, the model theoretically has no limitations, compatible with any UniMod PTM, and no restrictions of sequence length or precursor charges.[16] Similarly, DeepLC has no apparent limitations on PTMs and supports sequences up to 60 residues in length.[33] Chronologer is also expected to be quite robust with regard to PTMs.[34] The Prosit 2019 model was trained only on tryptic peptides with fragment charges of +1 to +3, with modifications limited to cysteine carbamidomethylation and methionine oxidation, and peptides up to 30 residues long.[12] Support for PTMs and HLA peptide training was achieved in the 2024 Prosit model.[35] In evaluating the performance of these models, it was immediately apparent that the empirical data reveals differing retention times of peptides in relation to their charge state (**Fig. S8A**); however, none of the five models evaluated displayed any variation with respect to precursor charge (**Figs. S8B-F**).Taken together, this indicates these prediction models do not represent the true variability in retention time with respect to precursor charge state.

### Evaluation of Spectral Libraries

After understanding the limitations of the RT models, we sought to understand if perhaps intensity models and RT models were ideal to combine. Initial results indicated that some intensity models, for example AlphaPeptDeep appeared to be incompatible with their corresponding RT model (**Fig. 3A-C**). Naturally, this was an unexpected result, leading to conclusions that perhaps these models are rather incompatible with DIA-NN, our analysis software of choice. Moreover, with respect to Prosit, it appeared that the RT model used with the library had very little effect on the number of neuropeptide identifications returned. Finally, the MS2PIP intensity models proved to be most successful overall, though this was consistently restricted to the Chronologer and the DeepLC RT models (**Fig. 3D**). Of note, the HCD 2021 model variant of MS2PIP did outperform the Immuno HCD iteration, perhaps lending to the biological differences of HLA peptides from neuropeptides.

**Figure 3).**
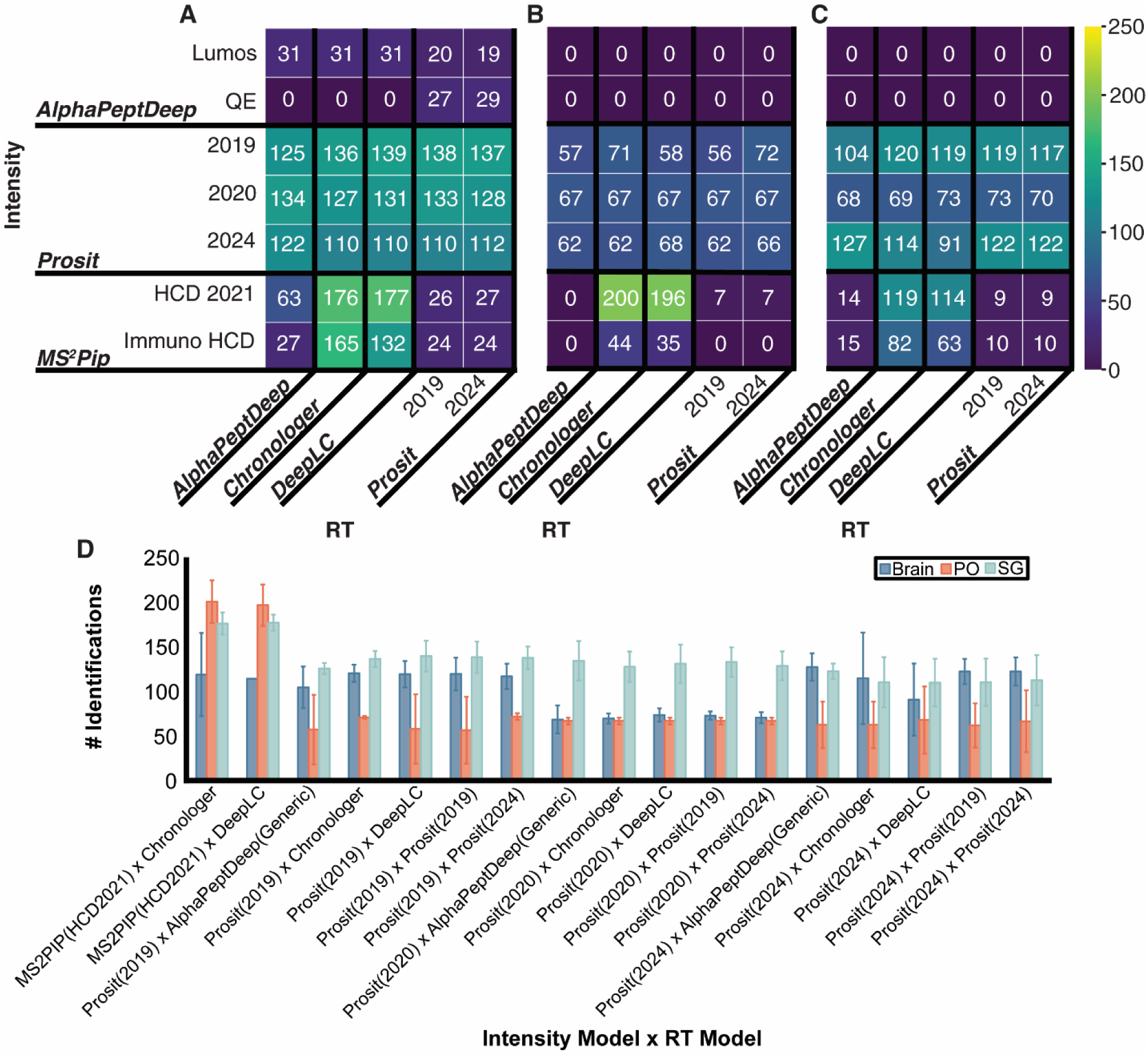
Evaluation of number of identifications returned with respect to intensity and retention time models used for **A)** sinus gland (SG), **B)** pericardial organ (PO), and **C)** brain tissue types. **D)** Global evaluation of number of identifications.

While we were pleased to find identifications from our predicted libraries in analysis of brain (**Fig. 4**), PO (**Fig. S9**), and SG (**Fig. S10**) tissue in DIA-NN, it was clear that the number of identifications did not scale with the size of the libraries. It should be noted that the empirical library represented about 50% of the know neuropeptidome, a common limitation in developing spectral libraries experimentally.[5] Despite this, in each scenario, it was evident that the empirical library yielded the highest number of identifications. Additionally, the largest subset of these results corresponded to the empirical library only. Despite this, it was clear that the new identifications were presented by the predicted models, for example 155 identifications were retrieved exclusively from the empirical library. Previously, it has been reported that library and library-free deconvolution approaches yield complementary results that together broaden the neuropeptidome; a similar trend is found in this work, where extending empirical libraries with predicted entries provides greater appreciation of the neuropeptides present.

**Figure 4:**
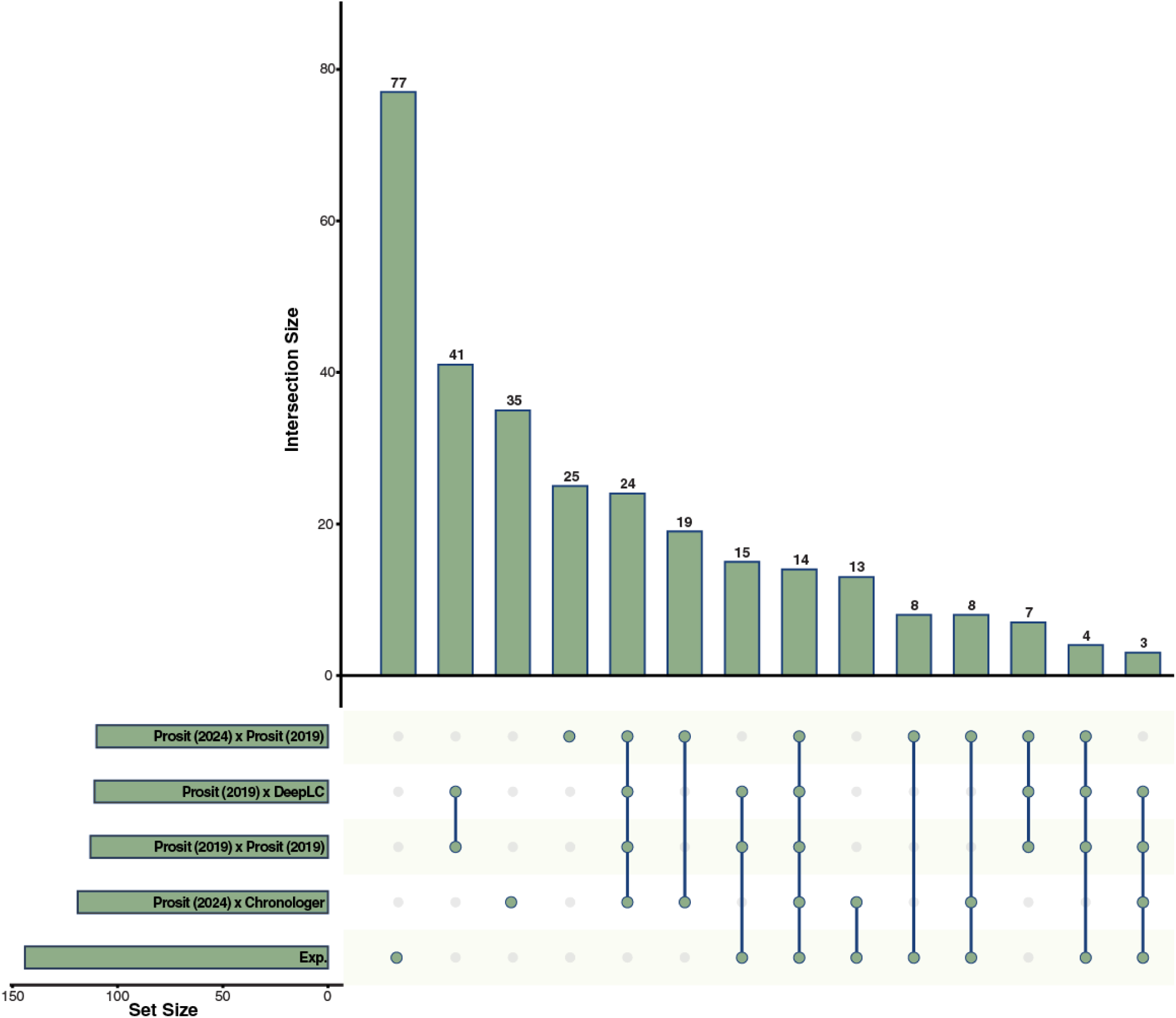
Intersection of all identifications produced via each model combination from brain tissue compared to the empirical (denoted as “exp”) spectral library.

Investigating further, we found that UniSpec’s key advantage is its inclusion of neutral loss fragments in spectra. Previously, we reported the high prevalence of neutral loss fragments in neuropeptide spectra.[10] Seemingly small perturbations to traditional HCD fragmentation or sample preparation can yield changes in neutral loss fragment presentation, for example with TMT-labeled peptides,[36, 37] or citrullinated peptides, stemming from the neutral loss of isocyanic acid.[38] Indeed, neutral losses have been deeply characterized in more targeted searches, for example in phosphopeptides,[39] or in more tailored fragmentation approaches, such as ETD,[40] but the frequency of these fragments appear under even the simplest of mass spectrometry conditions (*i.e.,* HCD, unmodified) has largely been overlooked. While the relationship is unclear at this point, it has been documented that 72% of b-ions within phosphopeptides yield neutral losses, compared to just 22% of y-ions, suggesting that b-ions are more likely to undergo neutral loss.[41] Of note, we have previously documented a continued observation that b-ions within endogenous peptides tend to be of higher abundance than their corresponding y-ions,[10] which could lend to the high frequency of neutral loss fragments in endogenous peptides. In profiling of our empirical library, it was clear that few peptides had an ion series that in total lacked any neutral loss ions. Indeed, the majority of peptides in our spectral library contained at least one fragment resulting from the loss of ammonium and water (**Fig. 5A**) If we evaluate all fragment ions within the library, approximately 45% of entries have no loss, and 26% are water loss and 28% are ammonium loss (**Fig. 5B**). With over half of all fragments arising from neutral loss, any accurate model must account for these ions in order to be relevant, particularly for neuropeptides.

**Figure 5:**
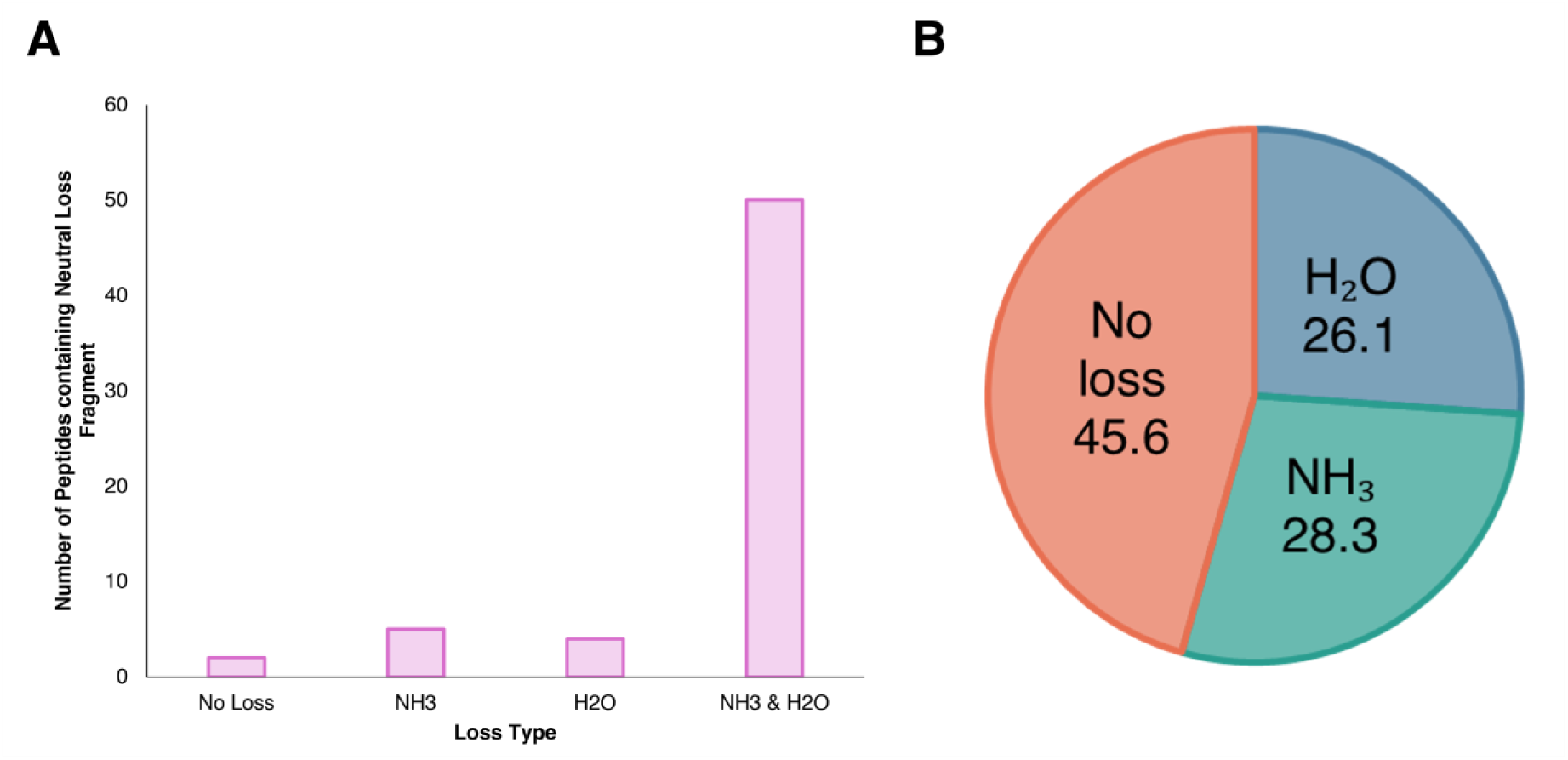
Evaluation of fragments with neutral loss(es) present in the empirical library corresponding to identifications only produced from the empirical library search. **A)** Number of peptides within this subset with fragments from neutral loss, **B)** percentage of neutral loss fragments in this subset.

### Conclusions

In this work, we provide a comprehensive analysis of popular spectral predictions models, a necessary task especially for sparse datasets where retraining or customization of a prediction model is infeasible. Here, we reveal the importance of neutral loss fragments in evaluation of neuropeptides and endogenous peptides, previously unevaluated. We also recognized that several models, even at exceptionally strict thresholds, are incapable of reaching a strong PCC to reflect consensus with the experimentally derived, empirical library with no modifications. This work highlights the unique strengths and pitfalls of each of the models, defining considerations required for building models for neuropeptides. In the future, a hybrid approach, integrating the qualities necessary for neuropeptide spectral prediction as derived here including consideration of a wide range of lengths and charge states, retention time prediction with respect to charge state, and inclusion of neutral loss ions would be beneficial in improving the effectiveness of identification neuropeptides from DIA analyses.

## Acknowledgements

This work was supported in part by National Institutes of Health (NIH) through grants R01DK071801, R01AG078794, and P41GM108538, and the Research Forward grant by University of Wisconsin - Madison Office of the Vice Chancellor for Research with funding from the Wisconsin Alumni Research Foundation (L.L.). The Orbitrap instruments were purchased through the support of an NIH shared instrument grant S10RR029531 and Office of the Vice Chancellor for Research and Graduate Education at the University of Wisconsin-Madison. L.F. was supported in part by the National Institute of General Medical Sciences of the National Institutes of Health under Award Number T32GM008505 (Chemistry–Biology Interface Training Program), the 2024 Eli Lilly and Company/ACS Analytical Graduate Fellowship, and a predoctoral fellowship supported by the NIH, under Ruth L. Kirschstein National Research Service Award (NRSA) from the National Institutes of Health-General Medical Sciences F31GM156104. L.L. would like to acknowledge NIH grants R01AG052324, S10OD028473, and S10OD025084, as well as funding support from a Vilas Distinguished Achievement Professorship and Charles Melbourne Johnson Professorship with funding provided by the Wisconsin Alumni Research Foundation and University of Wisconsin-Madison School of Pharmacy.

## Data Availability Statement

All Python scripts used for analysis are freely available at https://github.com/lingjunli-research/NeuropeptideSpectralLibraryPrediction. All mass spectrometry data and results have been deposited to the ProteomeXchange Consortium via the MassIVE partner repository with the dataset identifier: MSV000100853.

