## Supporting Information for "Benchmarking Spectral Library Prediction Platforms for Neuropeptidomics Applications"

### Table of Contents

Supplemental Figures (located within this document)

- **Figure S1:** Empirical spectral library properties
- **Figure S2:** Model evaluation via precursor charge
- **Figure S3:** Model evaluation via PTM
- **Figure S4:** Instrument-specific model evaluation
- **Figure S5:** Comparison of predicted models to empirical
- **Figure S6:** Comparison of all models to each other
- **Figure S7:** Normalization of all models based on returned IDs
- **Figure S8:** Retention time model evaluation
- **Figure S9:** IDs returned in DIA evaluation of pericardial organ tissue
- **Figure S10:** IDs returned in DIA evaluation of sinus gland tissue

Supplemental Tables (located within this document)

- **Table S1:** Comparison of model input requirements

Supplemental files

- **Supplemental File 1:** Queried peptides

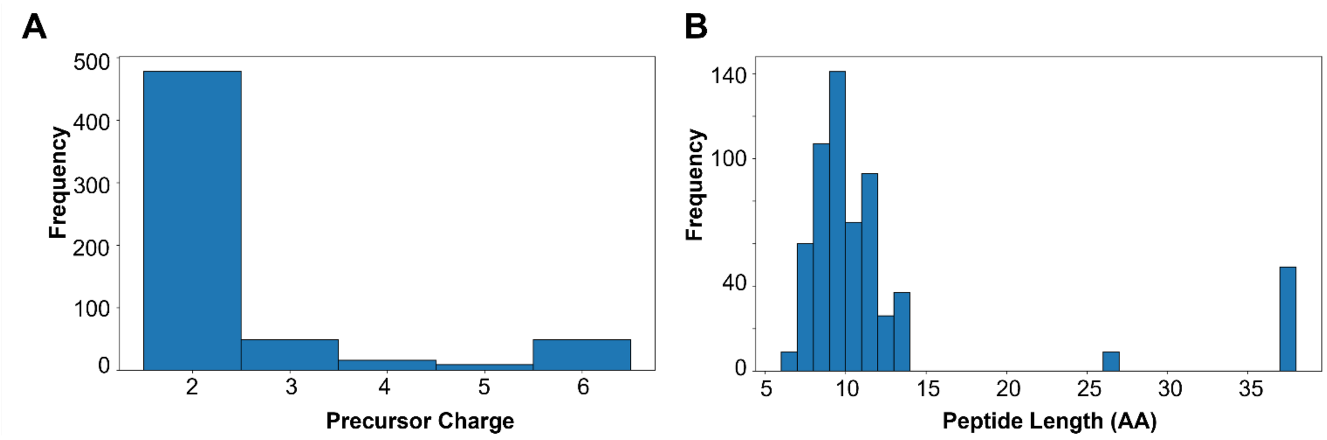


**Figure S1:** Properties of empirical spectral library. **A)** Length of all entries. **B)** Precursor charges of all entries.

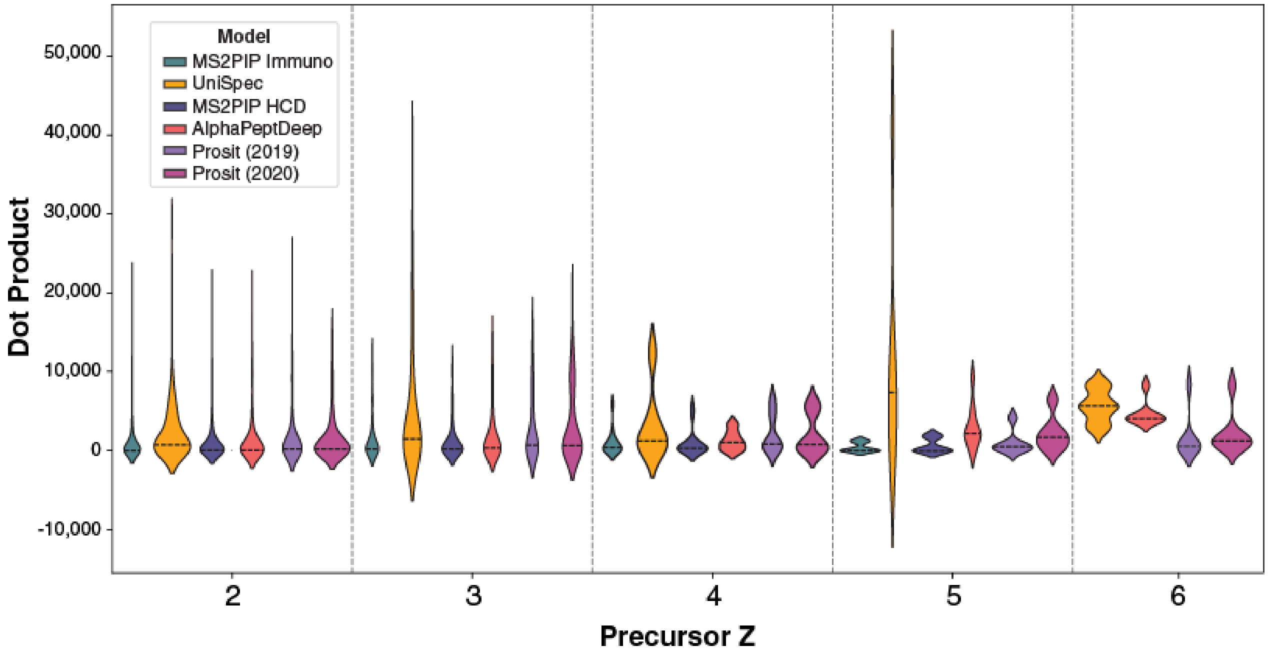


**Figure S2:** Corresponding dot product distribution with respect to precursor charge for each intensity model.

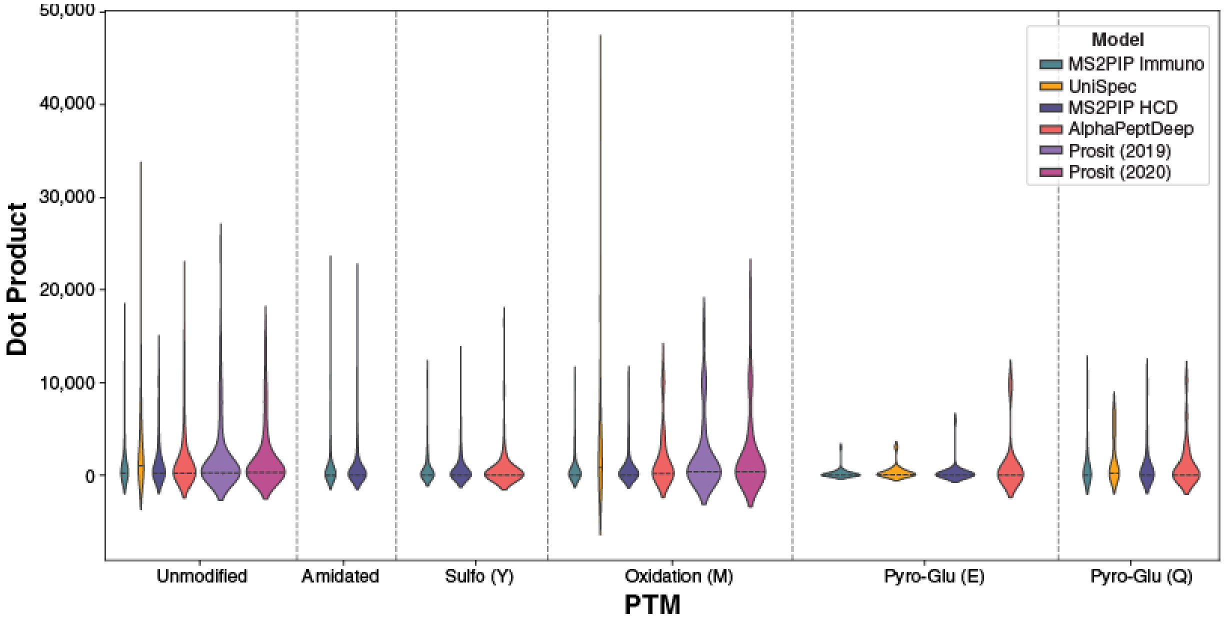


**Figure S3:** Corresponding dot product distribution with respect to the present post-translational modification for each intensity model.

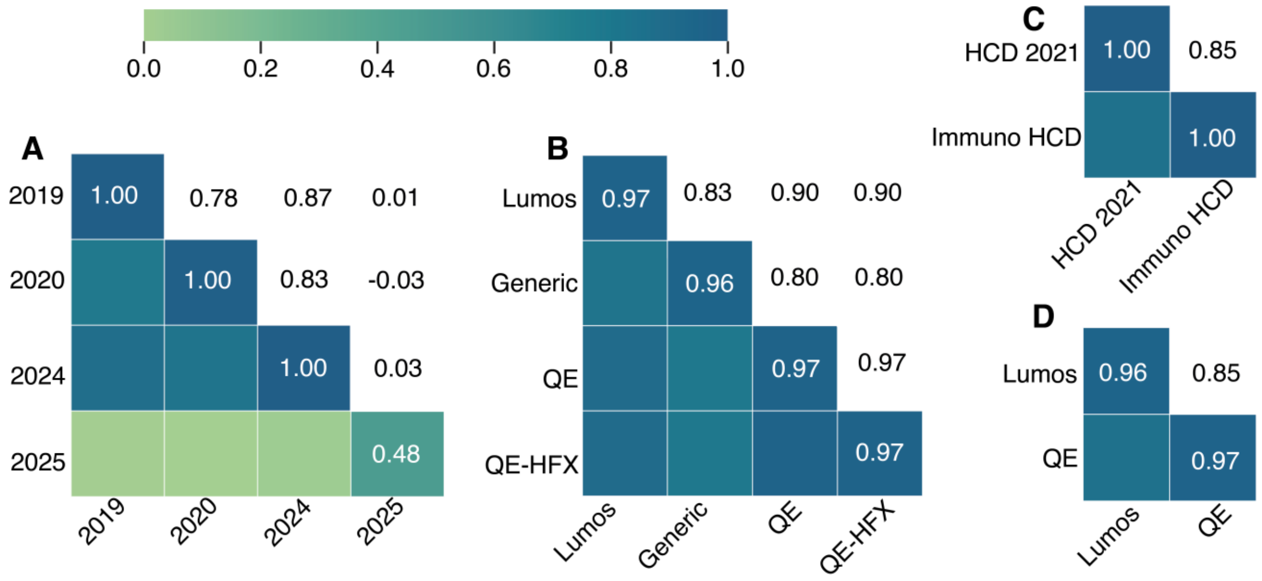


**Figure S4:** Pearson correlation coefficient of models released sequentially from the same research group. **A)** Prosit models from 2019, 2020, 2024, and 2025. **B)** UniSpec models generated for Thermo Lumos, QE, and QE-HFX Orbitrap instruments, as well as a generic model. **C)** MS2PIP models for HCD 2021 and immunopeptides HCD. **D)** AlphaPeptDeep models for the Lumos and QE Orbitrap instruments.

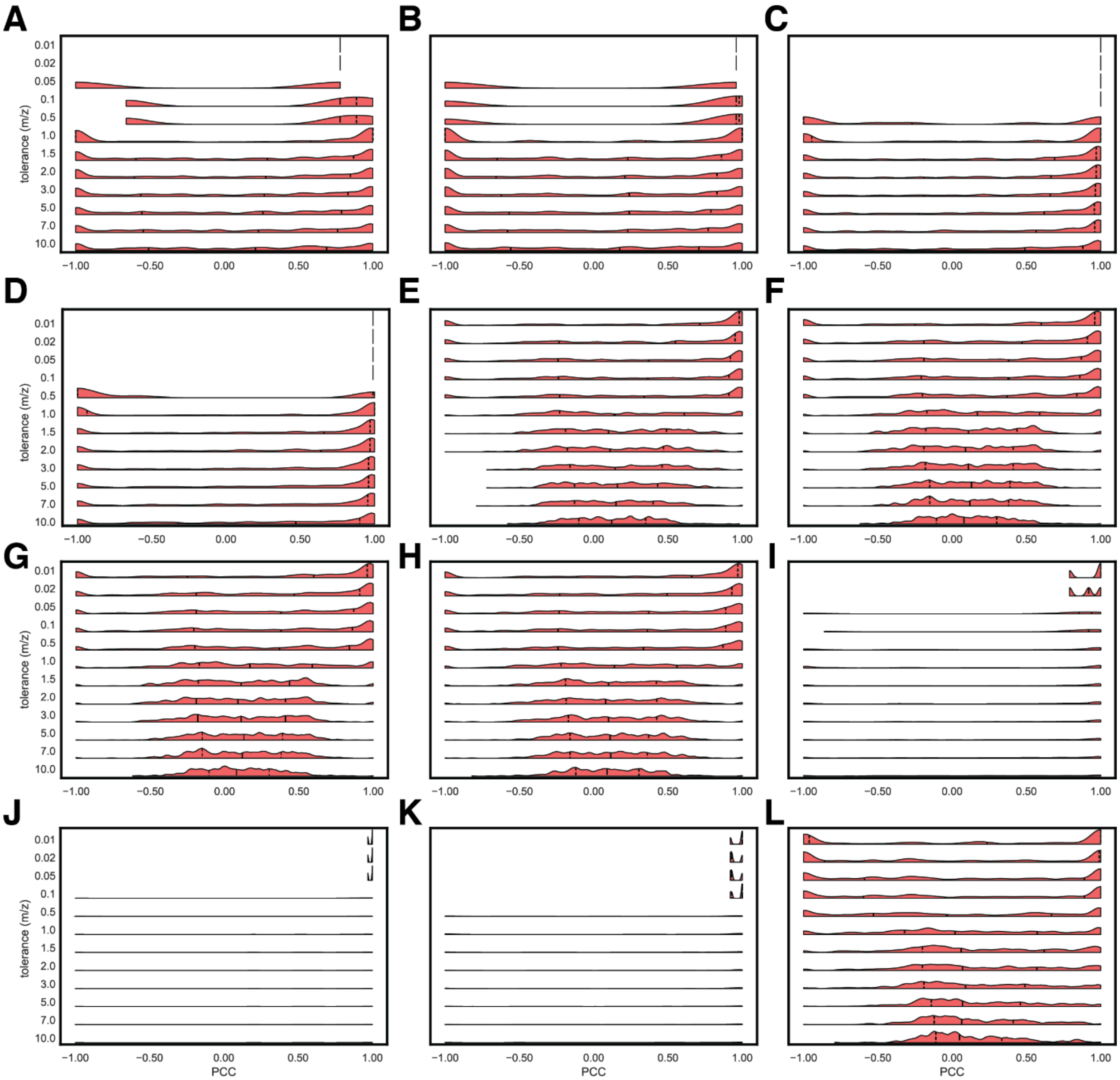


**Figure S5:** Sensitivity of models represented by PCC between the model and the empirical library. Models evaluated were **A)** MS2PIP HCD, **B)** MS2PIP Immuno, **C)** AlphaPeptDeep Lumos, **D)** AlphaPeptDeep QE, **E)** UniSpec Lumos, **F)** UniSpec QE, **G)** UniSpec QE-HFX, **H)** UniSpec Generic, **I)** Prosit 2019, **J)** Prosit 2020, **K)** Prosit 2024, **L)** Prosit 2025.

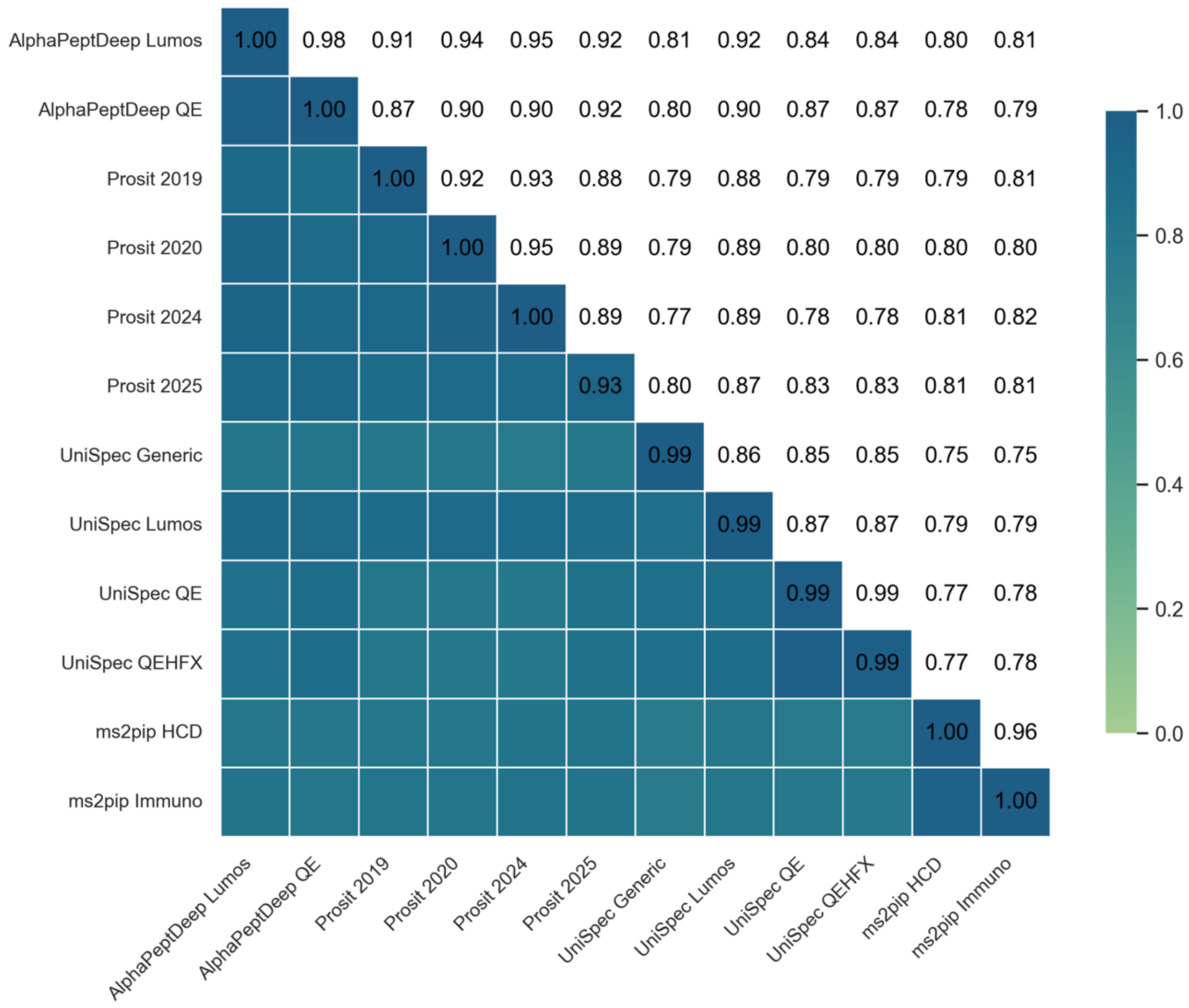


**Figure S6:** PCC of each intensity model with all other models without any PTMs.

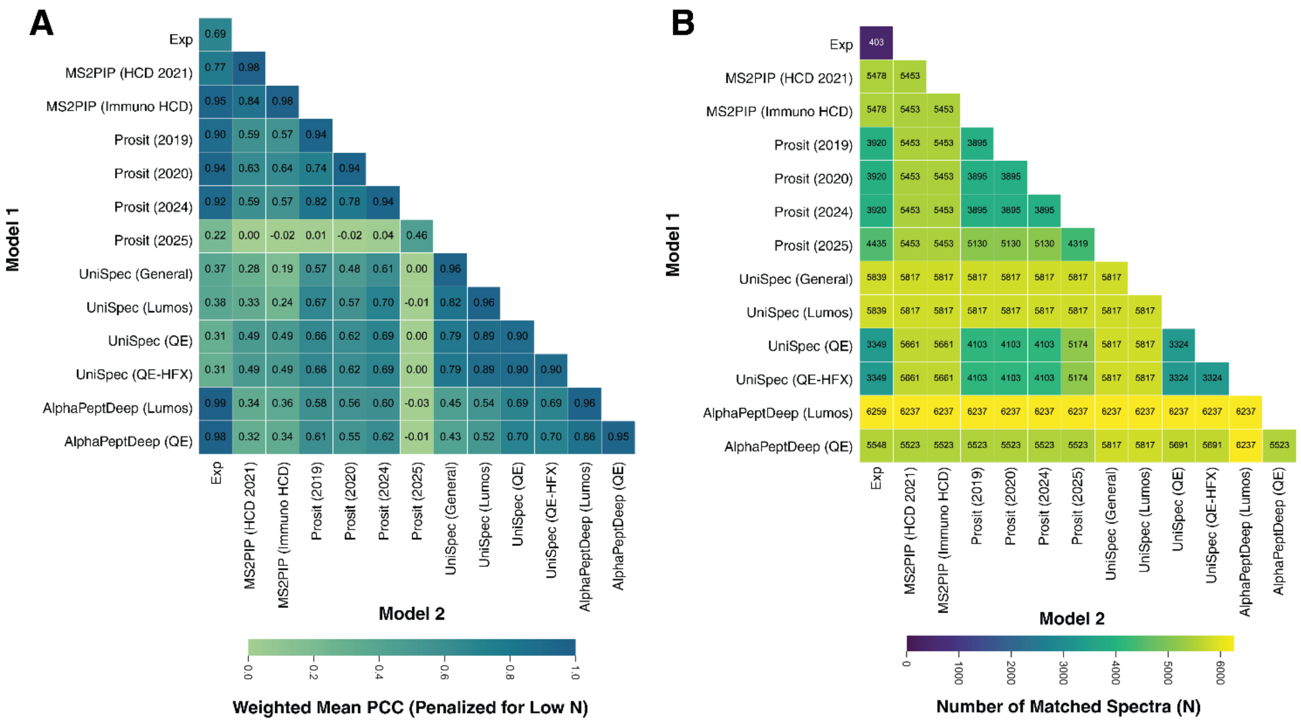


**Figure S7: A)** Weighted mean PCC for data illustrated in Figure 2, normalized to the number of peptide predictions produced by each model. **B)** Number of peptides used in weighting calculation. Exp refers to empirical spectral library.

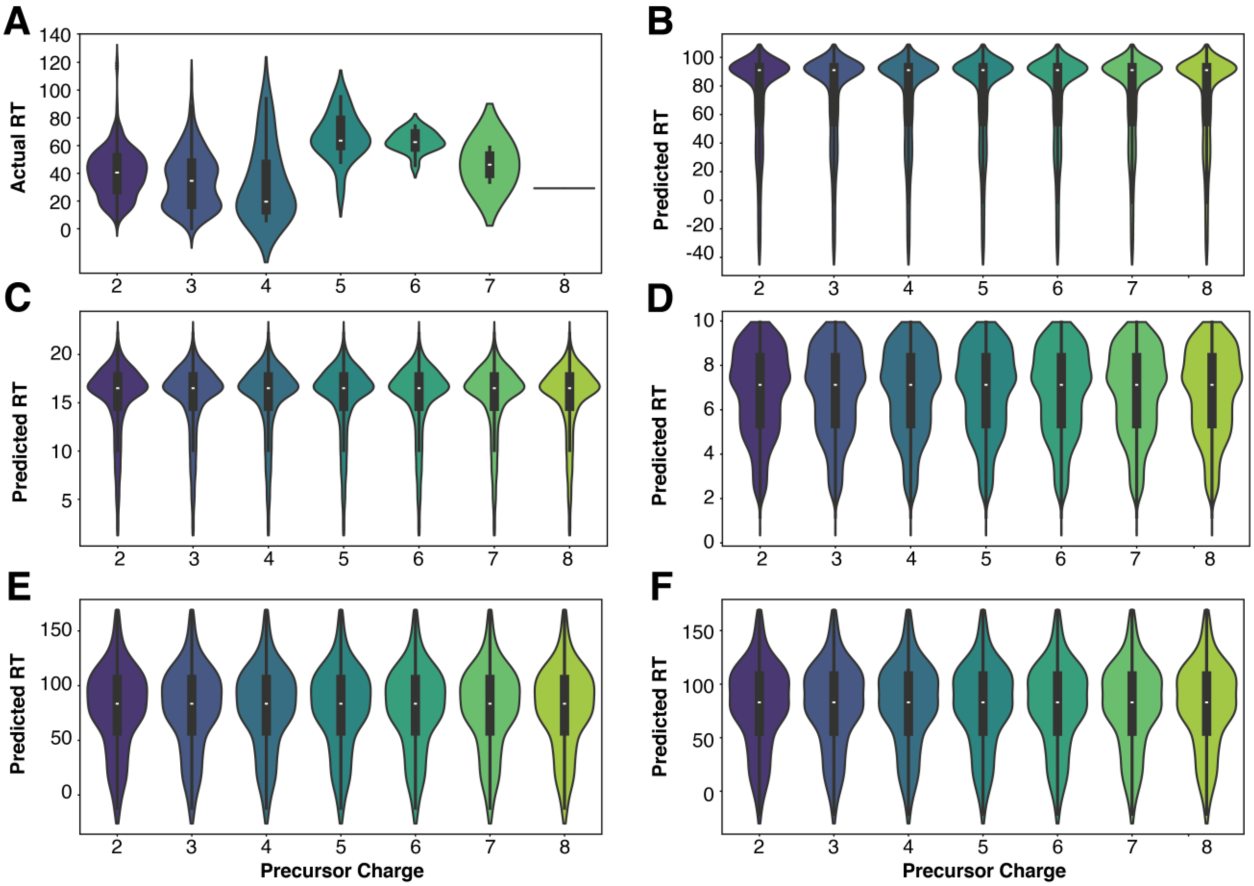


**Figure S8:** Evaluation of retention time models with respect to precursor charge. **A)** Empirical library. Models evaluated were **B)** AlphaPeptDeep, **C)** Chronologer, **D)** DeepLC, **E)** Prosit 2019, **F)** Prosit 2024.

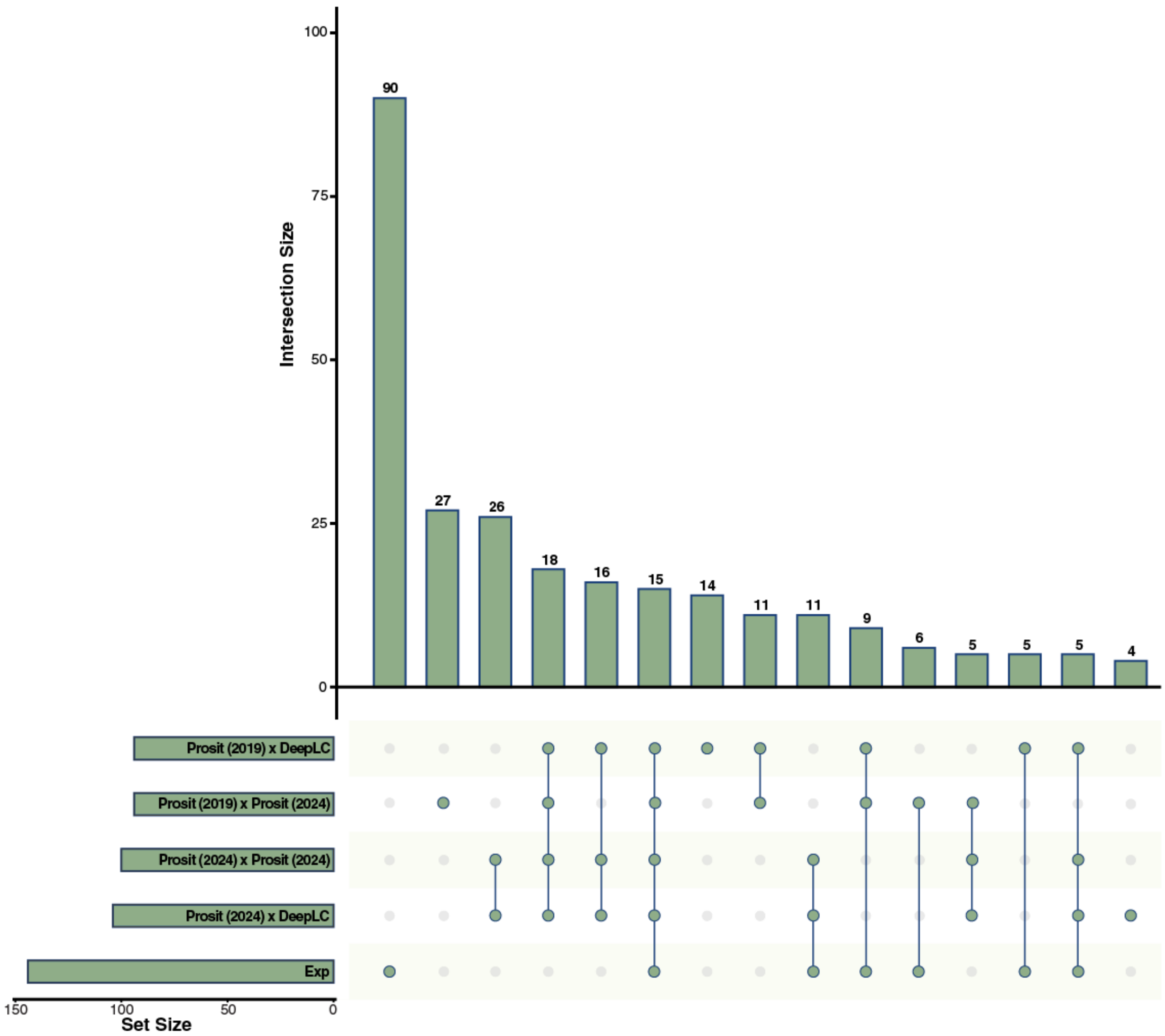


**Figure S9:** Intersection of all identifications produced via each model combination from pericardial organ tissue compared to the empirical (denoted as "exp") spectral library.

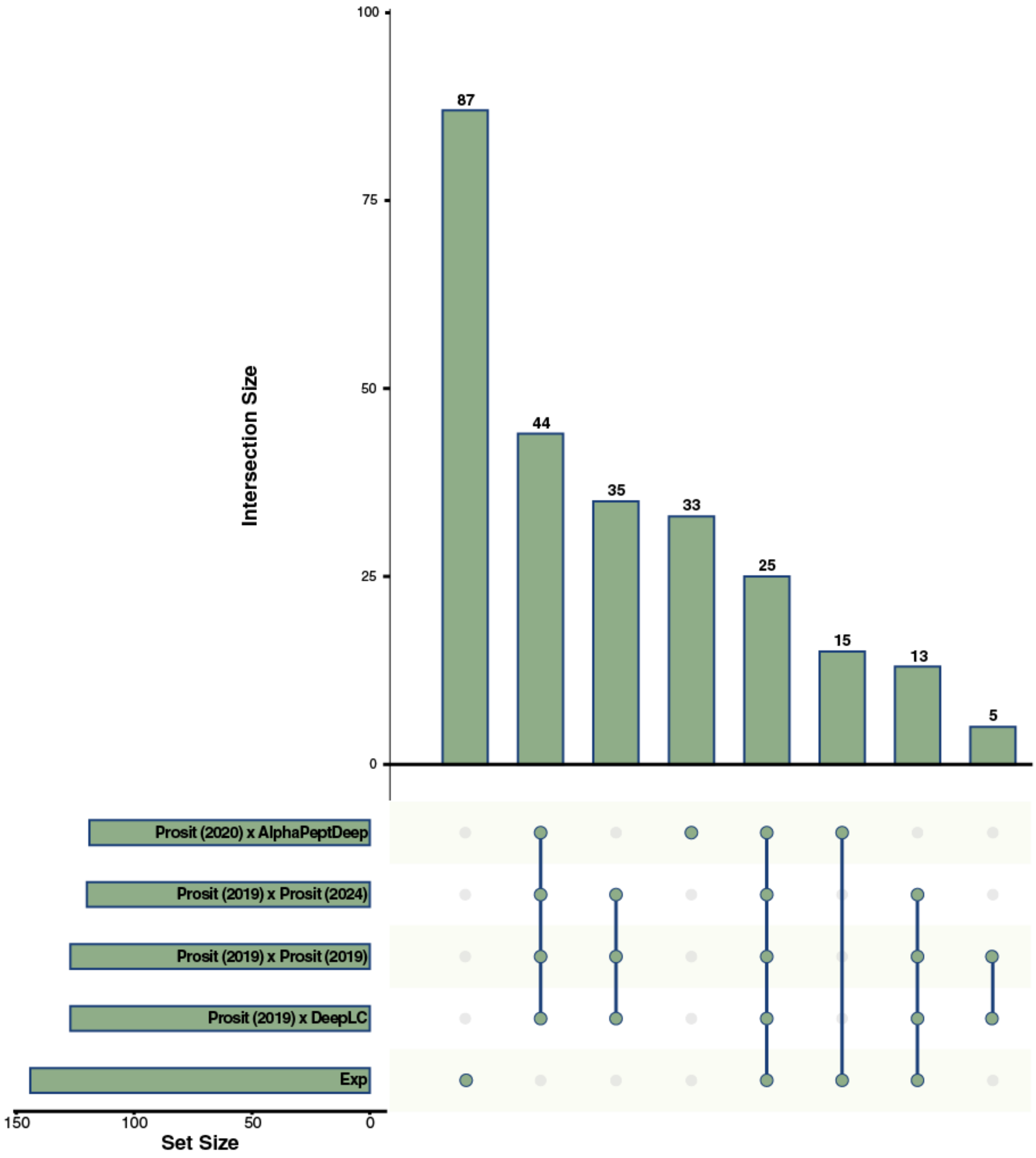


**Figure S10:** Intersection of all identifications produced via each model combination from sinus gland tissue compared to the empirical (denoted as "exp") spectral library.

**Table S1:** Comparison of each intensity model and their requirements for input peptides.

| | Training Data | | PTMs? | Max length |
|---|---|---|---|---|
| | Tryptic | Non-tryptic | | |
| Prosit 2019 | ✓ | ✗ | ✗ | 30 |
| Prosit 2020 | ✓ | ✓ | ✗ | 30 |
| Prosit 2024 | ✓ | ✓ | ✗ | 30 |
| UniSpec | ✓ | ✗ | ✓ | 40 |
| MS2PIP 2021 | ✓ | ✗ | Limited | 30 |
| MS2PIP Immuno | ✓ | ✓ | Limited | 30 |
| AlphaPeptDeep | ✓ | ✓ | Limited | N/A |
